# Functional mRNA delivery to hematopoietic stem and progenitor cells *in vivo*

**DOI:** 10.1101/2022.12.15.520650

**Authors:** David Alvarez, Guillemette Masse-Ranson, Saikiran K. Sedimbi, Phylicia Wisti, Lisa Rodriguez, Jordan Santana, Taylor Manning, Tim Towner, Ben Geilich, Cosmin Mihai, Ankita Mishra, Sushma Gurumurthy, Josh Frederick, Ulrich H. von Andrian, Jonathan Hoggatt, Melissa J. Moore, J. Rodrigo Mora

## Abstract

Gene correction of hematopoietic stem cells (HSC) is a promising therapeutic approach for multiple disorders. Current methods, however, require HSC collection from patients, gene correction during *ex vivo* culture, and re-infusion of corrected HSC into patients conditioned with chemotherapeutic agents. These approaches are complex, and the conditioning creates toxicities. We show that a lipid nanoparticle (LNP) can deliver mRNA encoding a reporter or a gene editing protein to HSC, with one injection transfecting ∼25% of mouse HSC, and repeated doses resulting in higher editing efficiencies. We also demonstrate LNP-driven *in vivo* mRNA delivery to HSC in non-human primates and humanized mice. These results demonstrate a translatable approach to deliver mRNA encoding therapeutic proteins, or gene correcting tools, to HSC that do not require cell culture or toxic conditioning.

**One-Sentence Summary:** LNP can deliver functional mRNA to mouse, non-human primate, and human HSC.

---

While mRNA-based medicines have been a promising concept for over two decades (*1*–*4*), only recently have they reached clinical validation with the unprecedentedly rapid development of high-efficacy vaccines against SARS-CoV-2 (*5*–*9*) as well as other infectious agents (*10*).

Clinical application of therapeutic mRNA beyond infectious diseases is quickly advancing in the oncology, autoimmune, and rare disease space (*11, 12*). Importantly, use of mRNA to deliver gene correction enzymes extends its potential to cure inherited and acquired genetic diseases (*13*).

Genetic defects in hematopoietic stem cells (HSC) cause many devastating life-threatening diseases with a wide spectrum of clinical manifestations, ranging from severe anemia and immunodeficiency to fulminant very-early-onset autoimmunity and inflammation (*14*). Success, however, relies on finding rare suitable histocompatible donors, and the procedure is often associated with significant morbidity and mortality due to opportunistic infections resulting from the preparatory conditioning regimen or graft-versus-host disease (GVHD) (*15*–*18*). A promising alternative approach is to correct the genetic defect in the patient’s own HSC and deliver the corrected HSC back to the patient. Currently, over 100 ongoing clinical trials are testing *ex vivo* HSC gene correction using a variety of approaches, including viral delivery of a new gene copy or genome editing approaches with zinc finger nucleases and Cas9 (*19*–*22*). While *ex vivo* gene correction does not require finding a donor, and may prevent GVHD, the approach is still limited by the need to successfully acquire high numbers of quality HSC for *ex vivo* manipulation.

Moreover, transplantation of the altered HSC can also result in many of the same morbidities associated with preparatory conditioning (*23*).

Use of LNP to deliver mRNA encoding either therapeutic proteins (*24*) or gene editing tools (*13*) to HSC *in vivo* would avoid the need for HSC collection, *ex vivo* culture, and toxic patient conditioning. Here we show that an LNP can efficiently deliver an mRNA encoding either a reporter or Cre recombinase to hematopoietic stem and progenitor cells (HSPC), including HSC, *in vivo*. We first assessed whether we can use LNP to reach HSPC in bone marrow (BM) by systemically administering an LNP comprising an ionizable aminolipid, distearoylphosphatidylcholine (DSPC), polyethylene glycol-(PEG) lipid, and cholesterol, plus a neutral phospholipid 1,2-dioleoyl-sn-glycero-3-phosphoethanolamine (DOPE) covalently-tagged to Atto647 (Atto647-LNP). After a single intravenous (i.v.) administration in mice we detected Atto647^+^ nucleated hematopoietic cells in blood, liver, spleen, and BM (Fig. 1A-C). At 5 min post LNP injection, ∼20% of BM hematopoietic cells were Atto647^+^ (Fig. 1C), indicating positive association with the systemically administered LNP. To determine whether the fluorescent signal in BM cells was from cells within blood vessels, we compared mean fluorescence intensity (MFI) levels in circulating (Gr1^hi^) monocytes to monocytes recovered across other organs over time (Fig 1 D). Mean fluorescence intensity (MFI) of Atto647 signal in BM monocytes was lower compared to blood monocytes suggesting that most Atto647^+^ cells in BM were not intravascular. Since at steady state HSPC are mainly confined to the BM niche (*25*), we assessed Atto647-LNP association with HSPC in this compartment by focusing on lineage-negative, Sca-1^+^, c-Kit^+^ (LSK) cells, which includes most progenitors as well as multipotent and self-renewable CD150^+^CD48^-^ HSC (*26*). Within 5 min of injection, ∼40% of HSPC were Atto647^+^, but rapidly decreased to background levels by 6 hours (Fig. 1E), possibly due to Atto647 cleavage from DOPE, as previously described (*27*). We also visualized sorted BM LSK cells using imaging flow cytometry, revealing punctate fluorescent Atto647 signal suggestive of internalized LNP (Fig. 1F).

**Figure 1.**
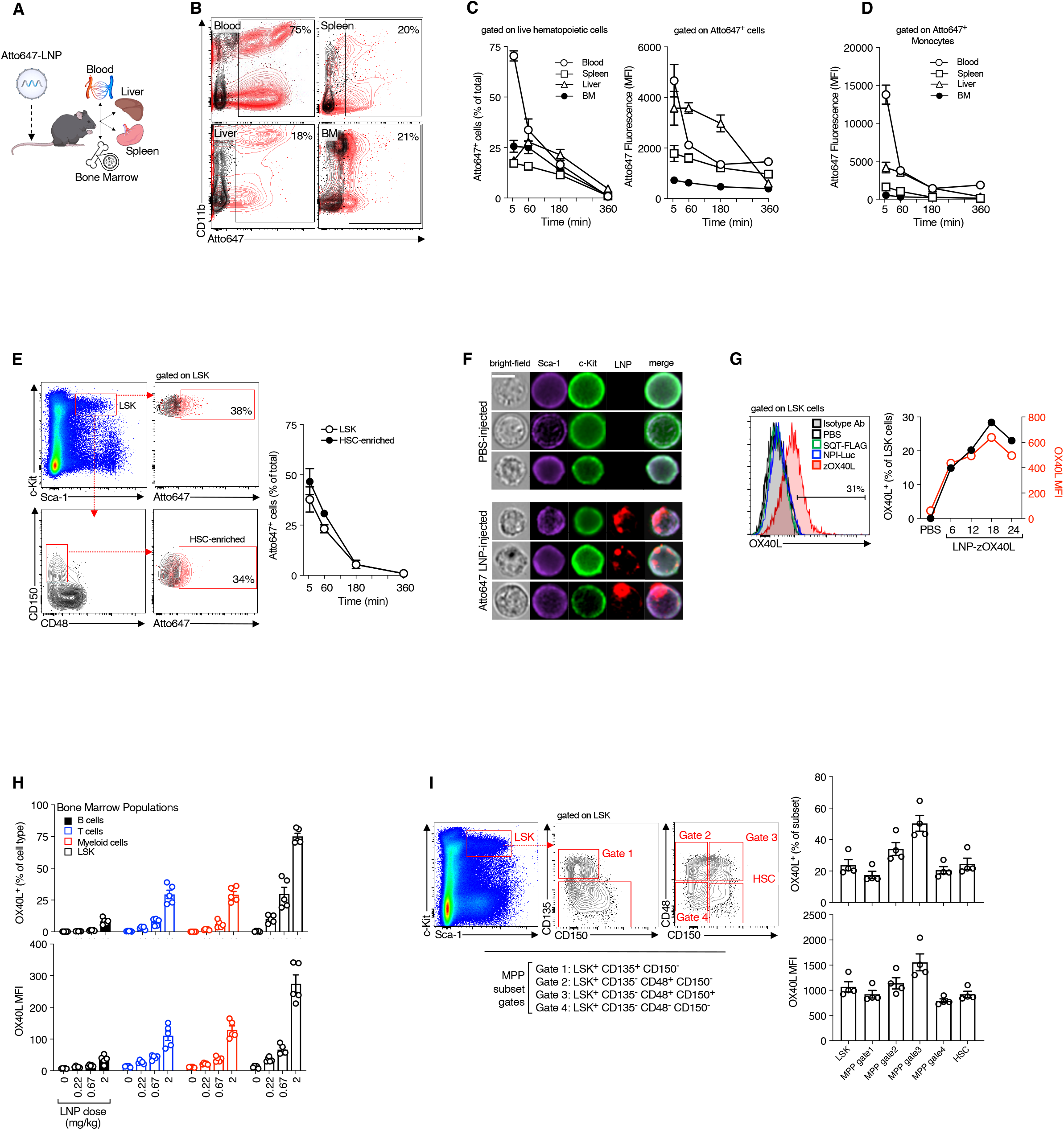
LNP-mRNA delivery to HSPC in the murine BM compartment. (**A**) Schematic illustrating fluorescent LNP distribution to different compartments at the cellular level following systemic i.v. delivery in mice (0.5 mg/kg). (**B**) Representative FACS plots showing Atto647^+^ detection in blood, liver, spleen, and BM cells. (**C**) Summary graphs quantifying frequency of hematopoietic Atto647^+^ cells (*left*) or Atto647 MFI (*right*). (**D**) Summary graph quantifying Atto647 MFI in Gr1^+^ monocytes. (**E**) Representative FACS plots depicting gating of Atto647 fluorescence in BM LSK cells and enriched CD150^+^CD48-HSC, and summary graph of Atto647 fluorescence in LSK and HSC gates. (**F**) Representative images of BM LSK cells post systemic injection of Atto647-LNP or PBS using imaging flow cytometry (ImageStream). Images depict Atto647 fluorescence (red) in enriched LSK cells stained for Sca-1 (purple) and c-Kit (green). Scale bar, 7 µm. (**G**). OX40L expression in LSK cells following delivery of LNP encapsulating mRNA for reporter zombie(z)-OX40L (LNP-zOX40L). Flow cytometry histograms (*left*) show OX40L staining in LSK cells from LNP-zOX40L-injected mice compared to isotype control antibody, or to mice injected with PBS or control LNP (SQT-FLAG, NPI-Luc). Summary graph (*right*) depicting kinetics of OX40L expression in BM LSK cells post i.v. LNP-zOX40L delivery. (**H**) OX40L expression in mouse LSK, lymphoid, or CD11b^+^ myeloid cells in BM upon administration of different LNP-zOX40L doses. Mean±SEM, n=5 mice/group. (**I**) Representative flow cytometry plots and summary bar graphs depicting frequency (%) and MFI of OX40L in different BM HSPC subsets upon a single i.v. LNP-zOX40L dose (0.5 mg/kg). Mean±SEM, n=5 mice/group.

Having established that Atto647-LNP can rapidly reach resident BM HSPC upon i.v. administration, we next assessed functional mRNA delivery by injecting LNP encapsulating mRNA encoding a non-signaling murine zombie(z)-OX40L (LNP-zOX40L), a reporter that can be readily detected by flow cytometry. While we could not detect OX40L in BM LSK cells in mice injected with either PBS or control LNP (SQT-FLAG, NPI-Luc), LSK cells from mice injected with LNP-zOX40L exhibited a ∼4-fold increase in OX40L MFI, with 18-32% of cells positive (Fig. 1G). Comparison among multiple cell populations in BM revealed greater OX40L expression (% and MFI) in LSK cells than in B cells, T cells, or CD11b^+^ myeloid cells at the two highest doses tested (Fig. 1H). To more comprehensively examine HSPC subsets for OX40L expression, we parsed LSK cells into multipotent progenitors (MPP; (*28*)) and HSC (Fig. 1I). At 24h post LNP delivery, approximately 18-50% of multipotent progenitors were OX40L, including HSC at ∼20%. Taken together, these data demonstrate that systemic LNP-mRNA administration in mice leads to rapid and efficient mRNA delivery to HSPC.

We next employed a mouse model that allows for lineage cell tracing. Ai14 mice contain a flox-STOP cassette sequence upstream of the gene encoding red fluorescent protein td-Tomato (tdT); cells in Ai14 mice remain non-fluorescent unless Cre-recombinase excises the flox-STOP cassette (Fig. 2A). Forty-eight hours after i.v. administration of LNP encapsulating mRNA encoding Cre-recombinase (LNP-cre), we observed robust tdT expression in 10-40% of HSPC subsets (Fig. 2B). We complemented the flow cytometry using functional *ex vivo* colony-forming unit (CFU) assays (*29*). Whereas colonies from vehicle-treated mice remained non-fluorescent, ∼30% of colonies from LNP-cre treated mice were tdT^+^ (Fig. 2C, D), consistent with the frequencies of tdT^+^ HSPC observed by flow cytometry. These data show that systemically administered LNP can deliver mRNA encoding a functional gene editing tool to BM HSPC *in vivo*.

**Figure 2.**
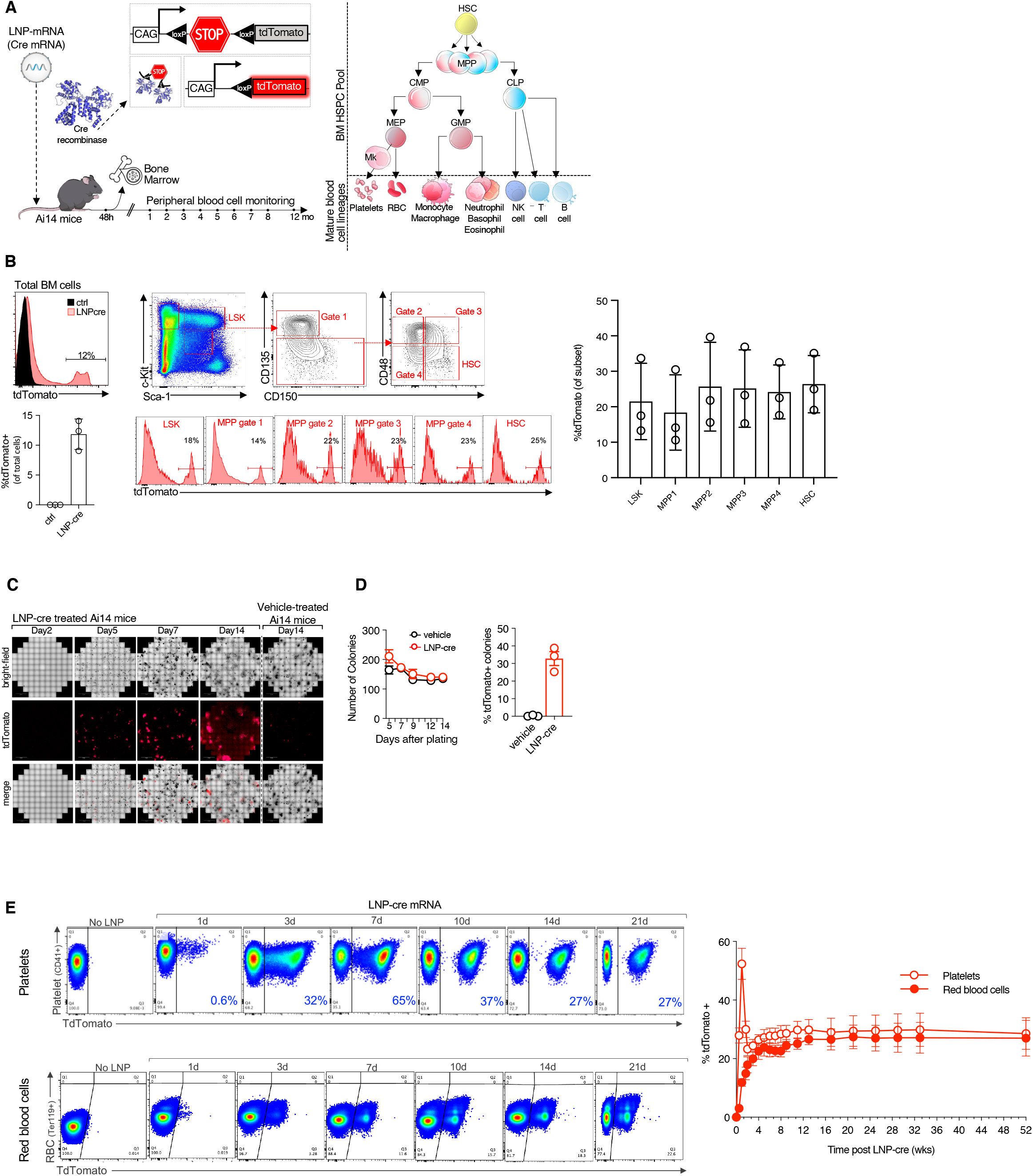
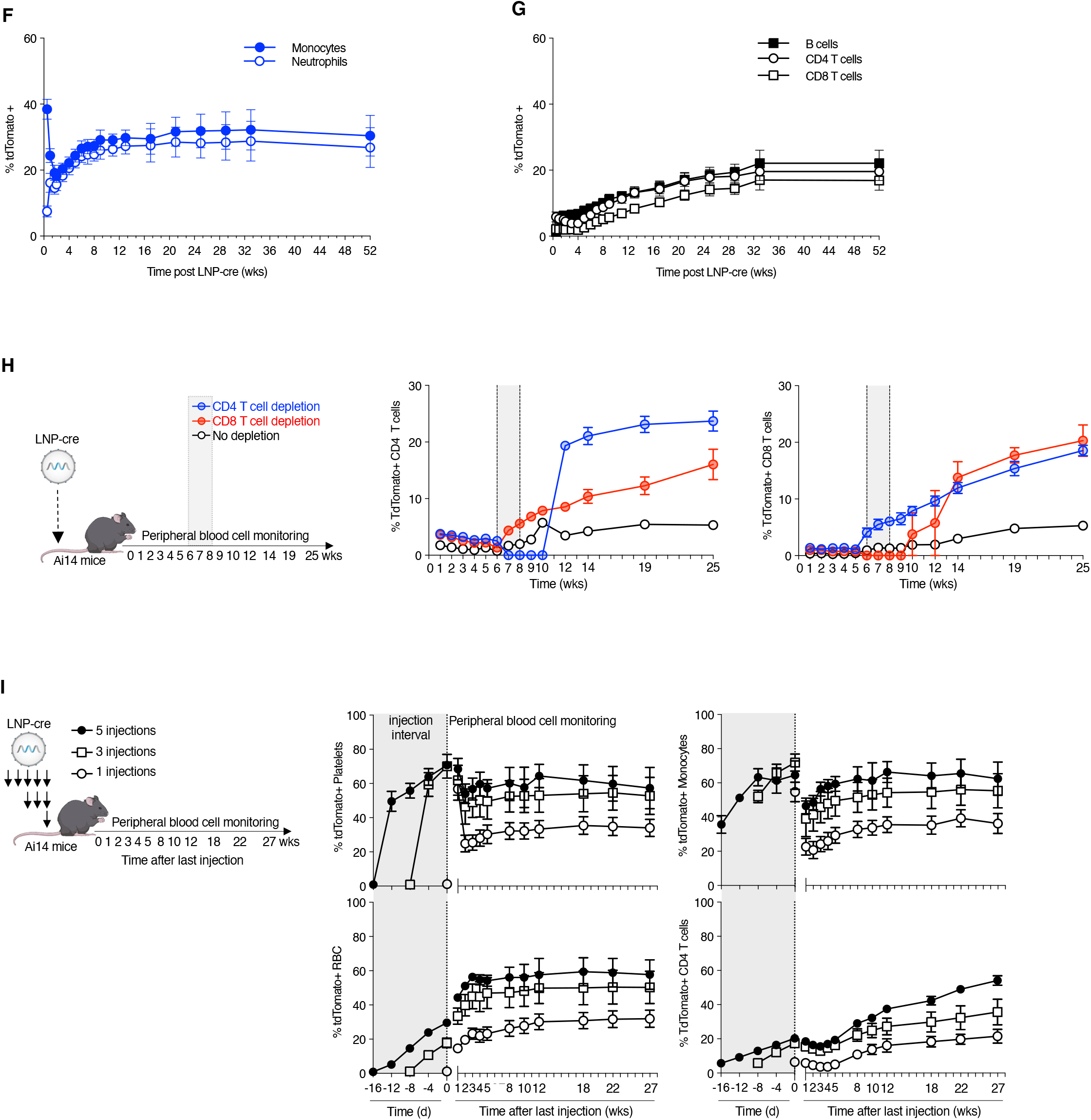
*In vivo* LNP-cre editing of BM HSPC in Ai14 mice leads to long-term myeloid and lymphoid labeling. (**A**) Schematic depicting *in vivo* editing of the Ai14-LoxP Stop cassette by LNP-cre mRNA delivery to track BM HSPC and mature cell lineages. (**B**) Induction of tdTomato (tdT) fluorescence in total BM cells and the HSPC pool following i.v. LNP-cre delivery or control (tris/sucrose vehicle). Representative flow cytometry gating of BM HSPC subsets, and representative histograms depicting tdT fluorescence. Graph shows frequency (%) of tdT^+^ among the HSPC subsets. Mean±SEM, n=3. (**C**) Colony forming unit (CFU) assay following *ex vivo* plating of BM cells harvested from Ai14 mice treated with LNP-cre or vehicle. Confocal microscopy showing brightfield (*top*), tdT fluorescence (*middle*), and merged (*bottom*) images at the indicated time points. (**D**) Summary graph depicting CFU counts at different time points, and frequency (%) of tdT^+^ colonies. Mean±SEM, n=3. **(E)** Lineage tracing analyses in peripheral blood following LNP-cre delivery in Ai14 mice. Representative flow cytometry plots and summary graphs depicting frequency (%) of tdT^+^ cells among total circulating platelets and RBC (**E**), monocytes and neutrophils (**F**), and CD4 T, CD8 T, and B cells (**G**) up to 12 months post LNP-cre delivery. (**H**) Schematic of transient lymphocyte depletion to assess effect on kinetics of tdT^+^ lymphocyte appearance in blood. Frequency of circulating tdT^+^ lymphocytes pre-and post CD4 (*left*) or CD8 (*right*) T cell depletion. (**I**) Multiple LNP-cre dosing leads to cumulative effect on HSPC editing Ai14 mice. Schematic of study design showing multiple LNP injections, with summary graphs depicting frequency (%) of tdT^+^ cells among circulating platelets and erythrocytes (*left*), and monocytes and CD4 T cells (*right*) up to 6 months post LNP-cre delivery. Dotted line at day 0 indicates the last injection performed for each of the three dosing groups. Each line represents a group (mean±SEM, n=3 mice/group). Shaded area in the plot depicts i.v. injection interval for LNP-cre, starting at day -16.

To establish the long-term impact of systemic LNP-cre delivery on hematopoietic lineages, we used flow cytometry to follow mature myeloid and lymphoid cells in blood over the course of 12 months. While at the earliest timepoint post injection (24 h) <1% of circulating platelets (CD41^+^) were tdT^+^, this fraction rapidly increased to ∼65% at 7d, and then plateaued to ∼30% after one month (Fig. 2E). Since platelets lack a nucleus and have a ∼5d lifespan in mice (*30*), the presence of tdT^+^ platelets in blood over 12 months indicates transfection of a long-lived upstream progenitor. The early spike at 7d was possibly due to transfection of mature megakaryocytes that were actively producing platelets at time 0. Ter119^+^ red blood cells (RBC) followed a different time course. Unlike platelets, there was no initial spike, but rather a slow increase in tdT^+^ RBC over time, plateauing at ∼25% after two months and continuing at that level for the entire 12 months (Fig. 2E). Like platelets, RBC also lack a nucleus. RBC, however, have a much longer lifespan of ∼40-50d in mice (*31, 32*), likely explaining the slower kinetics of approach to steady state.

Next, we examined nucleated myeloid cells including monocytes and neutrophils. At 24 hrs, the monocyte population exhibited a transient spike in tdT positivity, which subsided and then gradually approached a plateau of ∼30% tdT^+^ cells over 12 months (Fig. 2F). The initial spike in tdT^+^ monocytes is suggestive of direct LNP uptake by mature circulating monocytes with a lifespan of 7-10d (*33*). Similarly, the fraction of tdT^+^ neutrophils also plateaued at ∼25% over 12 months. These long-term data clearly indicate that the upstream progenitors of both monocytes and neutrophils were functionally transfected with LNP-cre.

Analysis of blood lymphoid populations in the same animals revealed slower kinetics, which reached ∼17-22% tdT^+^ cells by 12 months (Fig. 2G), consistent with the ∼2 month or ∼4-6 month lifespan of circulating B and T cells in mice, respectively (*34*). In fact, transient depletion of mature CD4 or CD8 T cells with specific antibodies during weeks 5-7 post LNP-cre administration led to rapid increase in the percentage of tdT^+^ CD4 and CD8 T cells to ∼20% of the circulating population (Fig. 2H). These data suggested that HSPC capable of differentiating to the full blood lineage spectrum were successfully edited with LNP-cre.

Having shown that a single LNP-cre administration could label 20-30% of the hematopoietic lineage, we next assessed the effect of multiple LNP-cre doses. We dosed Ai14 mice 1, 3, or 5 times, with injections spaced 4d apart. While mice receiving a single dose plateaued around 30% tdT^+^ platelets, RBC, and myeloid cells, those receiving 3 and 5 doses plateaued around 50% and 60% tdT^+^ cells, respectively (Fig. 2I, fig. S1). Lymphoid cells exhibited similar trends, with lower overall percentages due to their slower cellular turnover (Fig. 2I, fig. S1). Thus, multiple systemic LNP injections can transfect and edit at least 50% of total HSPC *in vivo*. Cumulative effects on gene editing suggest that a therapeutic threshold might be achieved by repeating LNP doses, even when using a relatively lower-efficiency gene-editing tool. Given the ability to assess the functional outcome of a corrected HSC and its progeny by sampling peripheral blood, patients could potentially receive customized numbers of treatments to achieve disease correction. By contrast, repetitive stem cell transplantation with the current *ex vivo* strategies is not practical or safe, particularly with existing conditioning regimens. Of note, for many diseases, correcting the majority of HSC *in vivo* is not necessary to achieve clinical benefit. For example, the morbidities associated with hemoglobinopathies like sickle cell anemia or beta thalassemia can be ameliorated with ∼20% normal HSC chimerism (*35*), with perhaps as low as 10% able to correct disease in some patients (*36*). In diseases that result in less competitive HSC, such as Fanconi anemia (*37, 38*), a smaller number of corrected clones may steadily increase in the patient, leading to therapeutic benefit. Our results with the current LNP in mice achieving 20% Cre-mediated editing after a single LNP dose, with up to 50% after 3 doses, suggests that therapeutic *in vivo* correction of a number of diseases is achievable and warrants further exploration.

Long-term hematopoietic lineage tracing in Ai14 mice suggested that LNP-cre transfect either HSC or long-lived progenitors. To formally establish whether systemic LNP-cre could transfect and edit HSC, we performed serial primary and secondary BM transplantations into lethally irradiated mice (Fig. 3A). Three Ai14 donor mice received a single i.v. dose of LNP-cre and, consistent with previous experiments, the frequency of tdT^+^ was ∼20% in blood non-lymphoid cells and ∼5% in lymphoid cells after 5 weeks (Fig. 3B-D, left panels), with 20-27% tdT^+^ BM LSK cells in donor Ai14 mice (fig. S2). BM cells were then transplanted into lethally irradiated CD45.1^+^ congenic (non Ai14) primary recipients (5 recipients per donor = 15 mice). In all primary recipients, the fractions of circulating tdT^+^ blood cell populations at 8 weeks were comparable to their respective Ai14 donor (Fig. 3B-D, middle panels). At 8 weeks post primary transplantation, we selected from each cohort a primary recipient with tdT^+^ blood cell frequencies approaching the average to serve as a donor for transplantation into 3-5 secondary CD45.1^+^ recipients. BM LSK cells from these donors ranged from 29-38% tdT^+^ (fig. S2). In secondary recipients we observed tdT^+^ platelets, RBC, myeloid, and lymphoid cells (Fig. 3B-D, right panels), with individual animals exhibiting similar tdT^+^ cell frequencies across myeloid and lymphoid lineages. These data demonstrate that bona fide BM HSC were transfected and edited *in vivo* upon a single LNP-cre injection in Ai14 mice.

**Figure 3.**
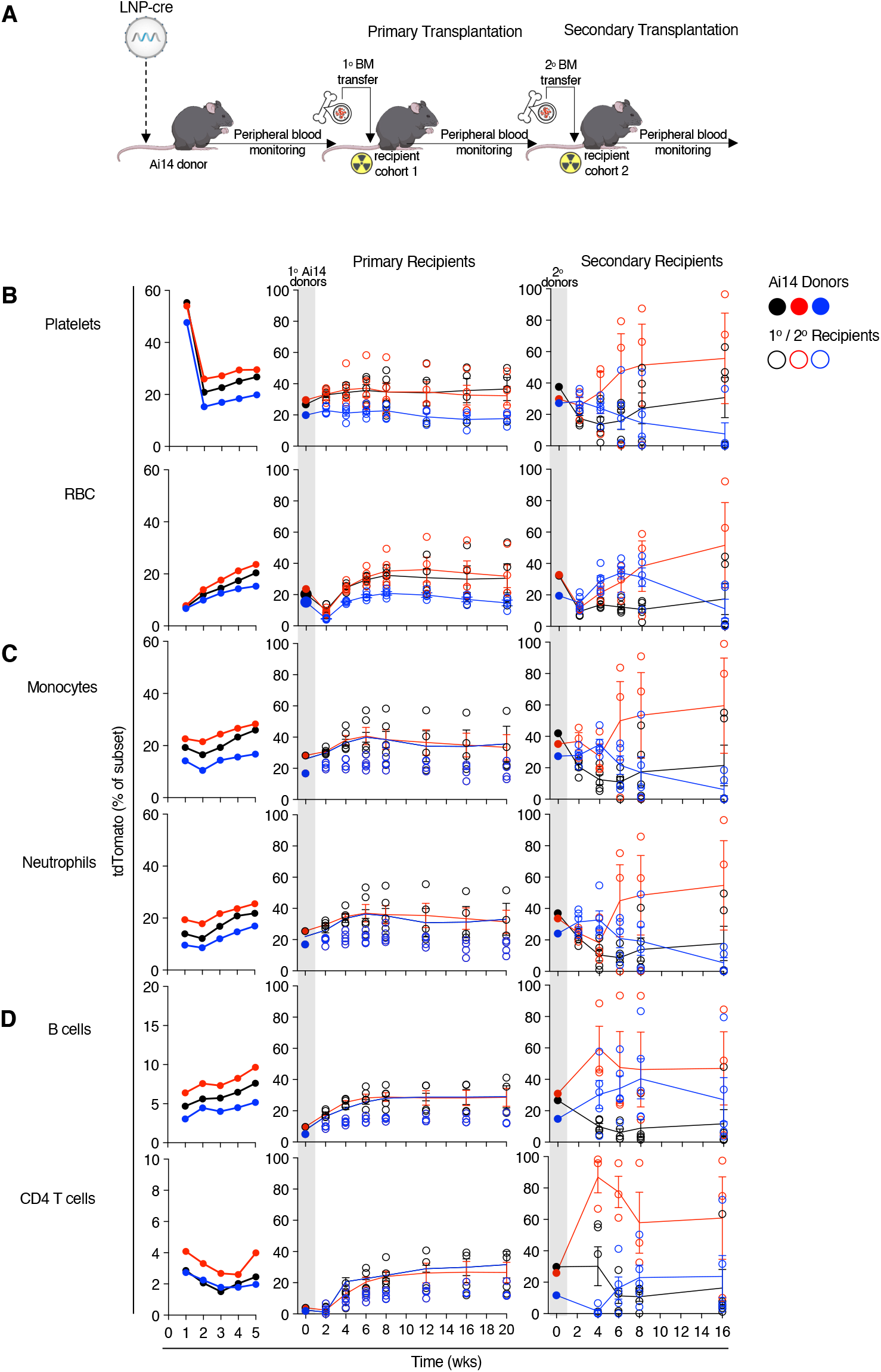
Full hematopoietic reconstitution upon serial BM transplantation provides evidence of *in vivo* LNP-cre delivery to bona fide HSC. **(A**) Schematic of serial primary and secondary BM transplantation into lethally irradiated mice. Frequency graphs depicting % tdT^+^ cells among (**B**) circulating platelets and RBC; (**C**) monocytes and neutrophils; (**D**) B cells and CD4 T cells, after primary and secondary BM transplantation. Grey shaded area and closed data points refer to blood profile of corresponding individual donor mice. Open data points refer to blood profile in the respective recipient mice. Immune cell subsets were gated on cells of CD45.2^+^/CD45.1^-^ Ai14 donor origin. (n=3 donors, n=10-15 mice recipients).

To assess whether our findings in mice translated to non-human primates, we dosed cynomolgus macaques with a single i.v. infusion of LNP-zOX40L and examined BM cells for mouse OX40L expression. We examined lineage-negative CD34^+^ BM HSPC, as well as the HSC-enriched CD90^+^c-Kit^+^CD45RA^-^CD123^-^ subset (Fig. 4A) (*39*). At 24h post LNP administration, OX40L was expressed in both lineage-negative CD34^+^ progenitors and the HSC-enriched subset. Both populations averaged ∼10% transfection, with some animals reaching over 20% (Fig. 4B), demonstrating that a single LNP administration can deliver functional mRNA to non-human primate HSPC *in vivo*.

**Figure 4.**
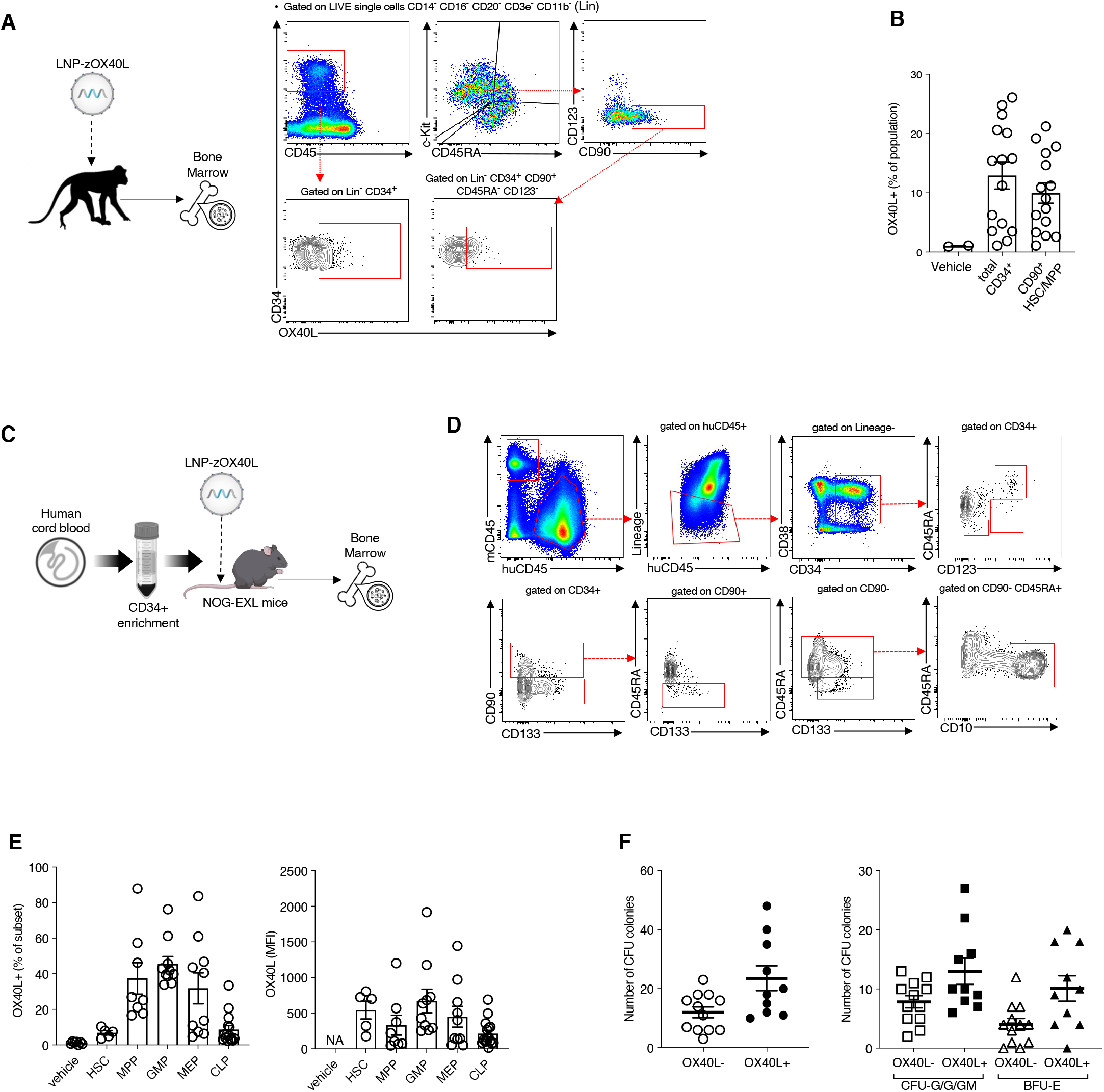
LNP-mRNA delivery to BM HSPC in non-human primates (NHP) and human HSPC in humanized mice. (**A**) Schematic of systemic LNP-zOX40L delivery to NHP *in vivo* and representative flow cytometry plots showing gating strategy for BM CD34^+^ HSPC and CD34^+^CD90^+^c-Kit^+^CD123^-^ HSC-enriched subset in NHP. (**B)** Summary graphs showing frequency (%) of BM cells expressing OX40L in total CD34^+^ HSPC and HSC-enriched subset. Each NHP received a single i.v. dose of LNP-zOX40L (0.3 mg/kg) and BM was collected 24h post injection. Mean±SEM, n=2 for NHP receiving vehicle, n=15 for NHP receiving LNP-zOX40L. (**C**) Schematic of LNP-zOX40L delivery to NOG-EXL humanized mice engrafted with human cord blood CD34^+^ cells and used 12-14 weeks post-engraftment. Mice were euthanized 24h after a single i.v. dose of LNP-zOX40L (0.5 mg/kg). (**D**) Representative flow cytometry plots depicting gating strategy used to analyze different human HSPC subsets in humanized mice. (**E**) Summary graphs showing frequency (%) of OX40L expression in human HSPC subsets. Data pooled from 3 independent experiments, with each dot representing one humanized mouse. Mean±SEM, n=7 for mice receiving vehicle, n=15 for mice receiving LNP-zOX40L. (**F**) Colony counts from CFU assays after plating FACS-sorted OX40L^+^ and OX40L^-^ human CD34^+^ HSPC from BM of humanized mice injected with LNP-zOX40L. Data pooled from 2 independent experiments; cells were sorted from n=5-6 humanized mice and plated in duplicate.

We then assessed LNP transfection ability of human HSPC *in vivo* in mice reconstituted with a human hematopoietic system (*40*). Immunodeficient NOG-EXL mice were xeno-engrafted with human umbilical cord CD34^+^ cells (Fig. 4C). At 24h post LNP-zOX40L injection, surface mouse OX40L was expressed in multiple different human HSPC subsets (Fig. 4D, E). CFU assays of FACS-sorted CD34^+^OX40L^-^ and CD34^+^OX40L^+^ cells demonstrated that both subsets were equally capable of producing myeloid and erythroid colonies (Fig. 4F), suggesting that human HSPC can be transfected *in vivo* without deleterious effect on hematopoietic progenitor potential.

Whereas *in vivo* hematopoietic gene editing has been achieved using viral vectors (*41, 42*), these vectors need to be customized for every target gene and they also elicit immunogenicity that often prevents repeated administration (*43, 44*). While our current work show that we can use LNP to deliver mRNA encoding Cre recombinase, this *in vivo* non-viral delivery approach is amenable to existing and emerging therapeutic gene editing tools (*45*–*48*). An additional challenge will be to develop LNP formulations capable of delivering Cas9 mRNA/gRNA for indications requiring gene correcting or replacing genes/exons (*49*–*51*). Lastly, in contrast to LNP delivering mRNA encoding therapeutic proteins, gene editing is potentially permanent.

Therefore, an important avenue of work will be to restrict the delivery or expression of gene editors as much as possible, which could be achieved, at least in part, by inserting tissue/cell-specific micro-RNA (miR) sites in the 3’ non-coding region of the mRNA encoding gene editors, such as Cas9. This strategy has been shown to be promising in immuno-oncology applications (*52*).

Our results demonstrate that an LNP formulation can consistently deliver mRNA to BM HSC/HSPC *in vivo* in mice and non-human primates, and to human HSC/HSPC in humanized mice. Long term lineage tracing and serial transplantation suggest that LNP can serve as vehicles for *in vivo* delivery of functional mRNA encoding therapeutic proteins or gene-editing tools to HSC/HSPC.

## Acknowledgements

We thank Megan Jesse, Ray Morrissette, Dan Roy, Jaclyn Milton, Michael Sokoler, Sarah Peterson, Claire Manuszak, Kimberyl Perez, David Easterhoff, Elena Shevtsova, Laureen Knowles, Graham MacLean, and Katy Olson for help with LNP formulations, *in vivo* studies, and helpful discussions.

## Disclosures

This study was funded by Moderna, Inc.

## Supplementary Figures

**Fig. S1.**
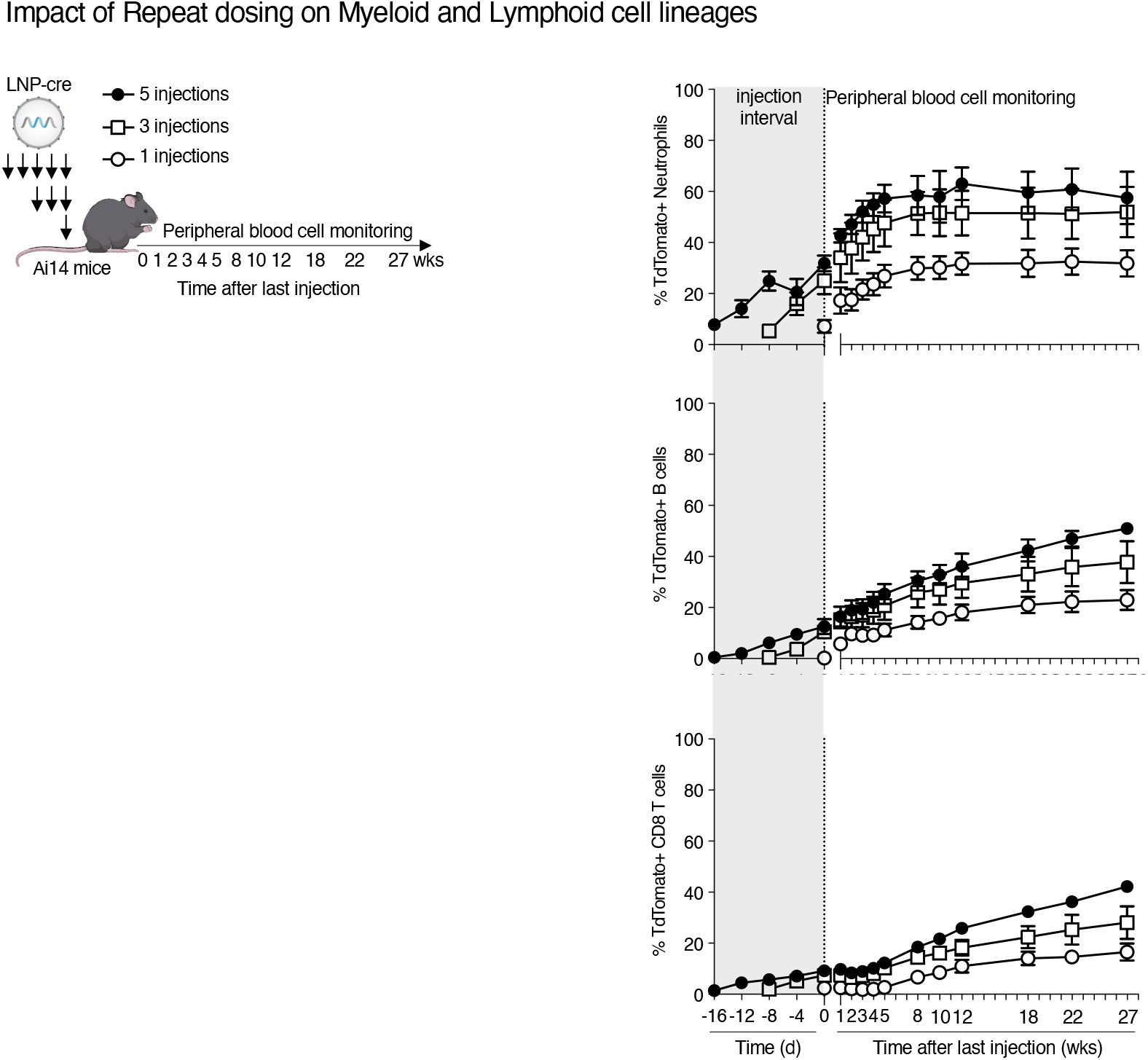
Multiple LNP-cre dosing leads to cumulative HSPC editing in Ai14 mice resulting in increased long-term myeloid and lymphoid cell labeling. Schematic of study design showing the effect of multiple LNP-cre injections (from main Fig. 2I) on the frequency (%) of tdT^+^ cells among circulating neutrophils (*top*), B cells (*middle*), and CD8 T cells (*bottom*) up to 27 weeks post LNP-cre delivery. Dotted line at Day 0 indicates the last injection performed for each of the three dosing groups, and the start of peripheral blood cell monitoring at weekly-to-monthly intervals. Data are mean ±SEM; n=3 mice per group.

**Fig. S2.**
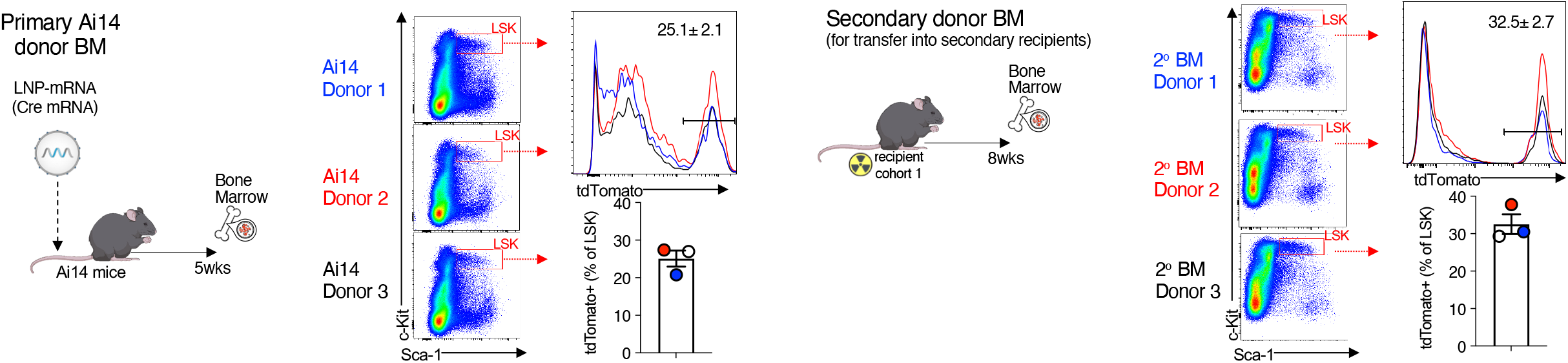
Frequency of tdT^+^ BM LSK cells in primary and secondary donors for serial transplantation. Schematic for primary Ai14 donor BM (l*eft*) and secondary donor BM (*right*) for use in serial transplantation experiments. For primary transplants, three Ai14 mice were euthanized 5 weeks post LNP-cre delivery (0.5 mg/kg) and the BM of each donor was transplanted into five CD45.1^+^ recipient mice (see Methods) and tracked for up to 20 weeks. From these primary recipients, three mice were selected to be donors for an additional cohort of CD45.1 recipient mice (secondary transplantation). For both primary Ai14 and secondary donors flow cytometric analysis was performed on BM. Flow cytometric pseudocolor dotplots and histograms depict frequency of tdT^+^ cells in the LSK gate for primary Ai14 donors (*left*) and secondary donors (*right*). Summary graphs reports mean ± SEM, n=3 donor mice.

## Materials and Methods

### Animal Studies

All mouse studies were approved by the Animal Care and Use Committee at Moderna and in accordance with NIH and US National Research Council guidelines on animal experiments and husbandry. Female and male C57Bl6 mice 6-8 weeks old were purchased from Charles River Laboratories. Homozygous female and male Ai14 (B6.Cg-Gt(ROSA)26Sortm14(CAG-tdTomato)Hze/J) and congenic CD45.1 (B6.SJL-Ptprca Pepcb/BoyJ) mice were purchased from Jackson Laboratory. Humanized mice were generated at Taconic Bioscience, using female NOD Cg-Prkdcscid Il2rgtm1SugTg [SV40/HTLV-IL3, CSF2] 10-7Jic/JicTac transgenic (NOG-EXL) mice engrafted with human umbilical cord blood CD34^+^. Humanized mice were used 11-18 weeks post engraftment, with at least 30% of human cell engraftment, and utilized for studies after 5d of acclimation. All mice were housed in specific pathogen-free (SPF) conditions on a 12h light-dark cycle, with food and water ad libitum.

Non-human primate (NHP) studies were conducted at Charles River Laboratories (Shrewsbury, MA) and approved by Charles River-MA Institutional Animal Care and Use Committee (IACUC) before conduct. Naïve male cynomolgus monkeys, 2–4 years old and weighing 2–5 kg, were housed in stainless steel, mesh floor cages, in a temperature-and humidity-controlled environment (18–29 °C and 30–70%, respectively), with an automatic 12-h/12-h dark/light cycle. NHP underwent tuberculin tests on arrival at the test facility. Animals were fed PMI Nutrition Certified Primate Chow No. 5048 twice daily and water ad libitum. To evaluate LNPmRNA delivery to BM HSPC, cynomolgus monkeys received a single dose of 0.3 mg/kg LNP-zOX40L or vehicle (20 mM Tris, 8% Sucrose, pH 7.5) by i.v. infusion. At 24h post administration, BM was flushed (humerus and femurs) and stored on ice for flow cytometry analysis.

### mRNA Synthesis and LNP Formulation

All mRNA constructs used in this study were generated *in vitro* by T7 RNA polymerase-mediated transcription from a linearized DNA template, with complete replacement of uridine by N1-methyl-pseudouridine as previously described (*53*).

LNP used in this study were manufactured via nanoprecipitation by mixing the ionizable lipid [(*53, 54*)], distearoylphosphatidylcholine (DSPC), cholesterol, and polyethylene glycol (PEG) lipid dissolved in ethanol with mRNA and diluted in sodium acetate buffer (pH 5.0). Formulations were concentrated as needed, passed through a 0.22 μm filter, and stored at 4 °C until use. Average LNP particle size was measured by dynamic light scattering (DLS) using the Wyatt DynaPro Plate Reader II (Santa Barbara, CA) and found to be <100nm. LNP-mRNA encapsulation efficiency (EE) was determined by the Quant-iT Ribogreen RNA assay (Life Technologies, Burlington, ON) as previously described [38]. All LNP formulations were quality controlled for particle size, mRNA encapsulation efficiency, and endotoxin levels prior to use for *in vivo*/*in vitro* studies.

### LNP administration

For most experiments, mice were dosed with LNP diluted in sterile PBS or tris/sucrose buffer by i.v. tail vein injection. To track fluorescent LNP distribution *in vivo* at the cellular level, mice were injected i.v. with a single dose of Atto647-LNP (0.5-1mg/kg) and euthanized at several time points post injection (5-10min, 1h, 3h, 6h). Organs/tissues (spleen, liver, BM, blood) were removed and immediately processed into single cell suspensions, stained and phenotyped by flow cytometry. To determine cellular LNP association/uptake, the frequency and MFI of Atto647^+^ cells were computed for each organ/tissue over time. To determine whether LNP can deliver mRNA to BM HSPC and other cells to mice *in vivo*, four-component LNP were formulated encapsulating different mRNA payloads (OX40L or Cre) and injected i.v. (for most experiments, dose range was 0.1-1.0 mg/kg). Isolated cells from blood, BM, and spleen were examined for reporter protein expression by flow cytometry (OX40L antibody staining or Cre-induced tdTomato fluorescence) compared to control mRNA constructs (SQT-FLAG or NPI-LUC) or vehicle (PBS or tris-sucrose)-treated mice. To determine the effect of multiple LNP injections on *in vivo* transfection of BM HSPC and downstream lineages, Ai14 mice were injected i.v. with 1, 3, or 5 injections of LNP-cre, 4d apart, and bled 24h after each injection to monitor induction of reporter expression (tdTomato fluorescence) in blood hematopoietic cells. Following the last injection, Ai14 mice were survival bled weekly, bi-weekly, or monthly for up to 12 months, and the frequency of tdT^+^ cells assessed over time.

### Tissue Collection and Flow Cytometry

For mouse tissue collection, LNP-treated, vehicle-treated, or control naïve mice were euthanized using CO2, and tissues/organs (blood, spleen, liver, and BM) were harvested and processed into single cell suspensions for flow cytometric analysis. Blood was directly collected into EDTA tubes by submandibular or cardiac puncture. A small aliquot was kept at RT and used for RBC and platelet staining, and the remainder blood sample subjected to RBC lysis and staining for myeloid and lineage cells and kept on ice. Spleen was cut into small pieces and plunged through a 70 µm cell strainer and centrifuged (450g, 5min). Resulting cell pellet was subjected to RBC lysis using ACK buffer for 1min, diluted with FACS-buffer, filtered, and centrifuged again before being counted and kept on ice for flow cytometry staining. For BM isolation, femur, tibia, and pelvic bones were harvested, flushed out with PBS, filtered through a 70um cell strainer and then centrifuged (300g, 10min). Resulting cell pellet was subjected to RBC lysis using ACK buffer for 1min, diluted with PBS, filtered, centrifuged, and resuspended in PBS or FACS-buffer and counted. For liver cell isolation, livers were harvested from mice following PBS perfusion (left ventricle), minced into small pieces and treated with enzymes (collagenase, DNase) for 30min at 37 °C under constant agitation. Liver suspension was then plunged through a 70 µm cell strainer and centrifuged (450g, 5min). Resulting cell pellet was subjected to RBC lysis using ACK buffer for 1min, diluted with FACS-buffer, filtered, and centrifuged again. Liver hematopoietic and non-hematopoietic (i.e. hepatocyte) fractions were not separated, and kept on ice for flow cytometry staining. For flow cytometry staining, cell suspensions were first stained for fixable viability dyes (Thermo Fisher Scientific) in PBS for 30min on ice. Cells were washed, then blocked with CD16/32 (clones 93, 2.4G2), FcγRIV (clone 9E9), and/or rat serum (2%; Stem Cell Technologies) for 5min on ice prior to adding fluorescent antibodies (BioLegend, BD, or Thermo Fisher Scientific) for cell surface staining. For RBC/platelet staining, cells were stained with CD41 (clone MWReg30), clone Ter119, CD71 (clone RI7217), CD44 (IM7), and CD45 (clone 30-F11). For peripheral blood leukocytes, cells were stained with CD41 and Ter119 (to exclude platelet-and RBC-leukocyte conjugates), CD11b (clone M1/70), CD115 (clone AFS98), Siglec-F (clone S17007L), Gr1 (clone RB6-8C5), CD4 (clone RM4-5 or GK1.5), CD8b (clone YTS156.7.7 or 53–6.7), B220 (clone RA3-6B2) or CD19 (clone 6D5). For spleen and liver, cells were stained with CD11b, CD115, Gr1, B220, CD45, and TCRb (clone H57-597). For mouse BM, cells were stained with lineage markers such as CD11b, CD4, CD8a, Ter119, Gr1, and B220, as well as stem cell markers CD117/c-Kit (clone 2B8), Sca-1 (clone D7), CD150 (clone TC15-12F12.2), CD34 (clone RAM34), CD48 (clone HM48.1), CD127 (clone A7R34), and CD135 (clone A2F10). For flow cytometry acquisition, cells were run on a BD flow cytometer (Fortessa or Fusion) and analyzed using FlowJo software (BD).

For human HSPC staining of BM cells, a panel including lineage markers (CD4 clone RPAT4, CD8a clone RPAT8, CD14 clone 63D3, CD19 clone 4G7, CD20 clone 2H7, CD11b clone LM2, CD56 clone 5.1H11, CD235a clone HI264) FITC, hCD45 APC-Cy7 (clone 2D1), CD34 BV650 (clone 561), CD38 AF700 (clone HIT2), CD90 PercpCy5.5 (clone 5E10), CD45RA APC (clone 5H9), CD123 BUV395 (clone 7G3), CD133 BV421 (clone S16016E), CD10 PE-TexasRed (clone HI10a), CD117(c-kit) BUV496 (clone YB5.B8), mCD45 PE (clone 30-F11), mOx40L PeCy7 (clone RM134L). Human HSPC subsets were defined as follows: HSC Long-term Hematopoietic stem cell (Lin-CD45^+^/dim CD34^+^ CD90^+^ CD133^+^ CD45RA-), MPP Multipotent progenitor (Lin-CD45^+^/dim CD34^+^ CD90-CD133^+^ CD45RA-), CLP Common Lymphoid Progenitor (Lin-CD45^+^/dim CD34^+^ CD90-CD10^+^ CD45RA^+^), GMP Granulocyte-Macrophage progenitor (Lin-CD45^+^/dim CD34^+^ CD45RA^+^ CD123^+^), MEP Megakaryocyte-Erythroid Progenitor (Lin-CD45^+^/dim CD34^+^ CD45RA-CD123-). Data were reported for each subset if >400 events were acquired. Lin-CD34^+^ zmOX40L^+^ and -cell sorting was performed using BD Fusion sorter.

For NHP tissue processing, BM was flushed (humerus and femurs) and lysed with ACK lysis buffer, then filtered through 70 µm cell strainers. NHP cells were stained for viability as detailed above, and then blocked with TruStain FcX (BioLegend) for 5min on ice prior to fluorescent antibody surface staining. NHP BM cells were stained with CD3e (clone SP34-2), CD4 (clone OKT4), CD20 (clone 2H7), CD11b (clone ICRF44), CD14 (clone M5E2), CD16 (clone 3G8), CD45 (clone D058-1283), CD45RA (clone 5H9), CD90 (clone 5E10), CD117/c-Kit (clone 104D2), CD123 (clone 7G3), and CD34 (clone 563).

### T Cell Depletion

For transient CD4^+^ T cell depletion(*55, 56*), mice were injected with 200 µg of anti-mouse CD4 antibody (clone GK1.5; BioXCell) in 200 µl of sterile PBS via the intraperitoneal (i.p.) route (3 doses, 4d apart). For CD8^+^ T cell depletion(*55, 56*), mice were injected with 200 µg of anti-mouse CD8b antibody (clone 53-5.8; BioXCell) via i.p. route (3 doses, 4d apart). After each antibody injection mice were survival bled 24h later and the degree of T cell depletion was determined using separate antibody clones. Following the third round of lymphocyte depletion, mice were survival bled weekly, bi-weekly, or monthly for up to 6 months, and the frequency of tdT^+^ cells among myeloid and lymphoid lineages was assessed and compared to pre-depletion levels or PBS-injected control mice.

### Colony Forming Unit (CFU) Assays

For mouse CFU assays, single cell suspensions from LNP-cre-or vehicle-treated Ai14 BM were obtained as described above and plated according to manufacturer recommendations (Stem Cell Technologies). Briefly, approximately 3×10^4^ total Ai14 BM cells were plated per well in MethoCult (M3434, Stem Cell Technologies) enriched with rmSCF, rmIL-3, rhIL-6, rhEPO, rh insulin, and human transferrin to support the optimal growth and differentiation of hematopoietic cells including erythroid progenitors (CFU-E/burst forming unit [BFU]-E), granulocyte and/or macrophage progenitor cells (CFU-GM), and multi-potent granulocyte, erythrocyte, macrophage, and megakaryocyte progenitors (CFU-GEMM). Plates were observed for colony formation and induction of tdTomato fluorescence by confocal microscopy and image analysis. For human CFU assays, 2×10^3^ Lin-CD34^+^ mOX40L^+^ and mOX40L^−^ were sorted and plated in MethoCult medium and CFU-G/M/GM and BFU-E (and marginally CFU-GEMM). Total colonies were numerated after 10d of culture. For NHP CFU assay, 1×10^3^ Lin-CD34^+^ CD45int CD45RA-CD123-CD90^+^ cells were sorted based on zmOX40L(±) reporter expression and plated in MethoCult (H4435, Stem Cell Technologies) enriched with rhSCF, rhIL-3, rhIL-6, rhEPO, rhG-CSF, and rhGM-CSF. Total colonies and colony subtypes were enumerated after 10 days.

### Primary and secondary BM transplantation

For primary BM transplantation, female Ai14 mice (CD45.2 background) were dosed with LNP-cre (0.5 mg/kg i.v.) and served as primary donors. At 5 weeks post LNP-cre, Ai14 donors were euthanized, and the femurs, tibiae, and pelvic bones were harvested, flushed out with PBS, filtered through a 70 µm cell strainer and then centrifuged (300g, 10min). Resulting cell pellet was subjected to RBC lysis using ACK buffer for 1min, diluted with PBS, filtered, centrifuged, and resuspended in PBS or FACS-buffer and counted. A fraction of BM cells were stained for flow cytometry to determine the frequency of tdTomato in donor cell fractions, and the remaining cells injected i.v. (5-7.5 ×10^6^/200 µl PBS) into lethally-irradiated congenic CD45.1 recipient female mice. Primary recipient mice were survival bled bi-weekly/monthly for up to 20 weeks, and analyzed for myeloid and lymphoid cell chimerism and frequency of tdTomato by flow cytometry. For the serial secondary BM transplantation, a cohort of primary BM recipients from the first BM transplantation were euthanized and served as secondary donors. Secondary donor BM was harvested as described above, and depleted of lineage-positive cells by magnetic columns (Stem cell technologies) to enrich for HSPC. Enriched BM cells were stained for flow cytometry to determine the frequency of tdTomato in secondary donor cell fractions, and the remaining cells injected i.v. (1 ×10^6^/200ul PBS) into lethally-irradiated congenic CD45.1 recipient female mice. Secondary recipient mice were survival bled bi-weekly/monthly for up to 16 weeks, and analyzed for myeloid and lymphoid cell chimerism and frequency of tdTomato by flow cytometry.

### Confocal Microscopy and Image Analysis

BM cells were harvested from Ai14 mice 48h post LNP-cre or vehicle administration, then plated *ex vivo* for up to 14d. BM cells were imaged 2, 5, 7, 9, 12, and 14d after *ex vivo* plating with an Opera Phenix high-content imaging spinning disk confocal system (Perkin Elmer) with a 5x air objective (NA 0.16). 97 fields of view were stitched together to form a global image of each well; n=2 for vehicle treated mice, n=3 for LNP-cre treated mice (Figure 2C). Brightfield images were obtained at 50% laser power with a 100ms exposure. TdTomato images were collected by illuminating with a 561nm laser at 100% power with a 400ms exposure. A z-stack of six optical sections spanning 25µm were acquired for both channels. Images were analyzed using Harmony software with PhenoLOGIC (PerkinElmer). Colonies were visible 5-14d after *ex vivo* plating. The total number of colonies per well were counted on each day, and the number of tdTomato+ colonies counted by analyzing images taken 14d after *ex vivo* plating.

### Imaging flow cytometry

Fluorescently labeled LNPs incorporating 0.1% ATTO 647 DOPE were used to quantify LNP association and uptake by HSPC. Mice treated with fluorescent (Atto 647) LNP were euthanized 5-10 mins, 1, 3 or 6 hours post i.v. injection. Blood, spleen, liver and BM were harvested, and single cell suspensions were stained for analysis by flow cytometry as described previously. To study direct LNP uptake and association by HSPC in the BM, stained BM cells were sorted (Aria Fusion cell sorter, BD Biosceinces) to obtain an enriched population of LSK cells (Lineage-, Sca-1^+^, c-Kit^+^) for further analysis by imaging flow cytometry using an ImageStreamX MkII (Amnis Corporation) equipped with 12 imaging channels to acquire fluorescent, bright field and dark field images for each cell. Enriched LSK cells were acquired with the following LASER power settings at slow speed and 60X magnification: 405nm: 150mW, 488nm: 200mW, 642nm: 150mW, 785nm: 2mW (Data acquisition software: INSPIRE). Acquired data (raw image files (RIF)) were analyzed using the data analysis software IDEAS. Single color controls were used to create a compensation matrix that was applied to all experimental files to correct for fluorescence spillover. Lineage negative single cells were gated in a bivariate plot of c-Kit and Sca-1 to identify double positive LSK cells. Single color and fluorescence overlayed images were generated to depict the cells containing Atto647 fluorescence.

## Notes

### Competing Interest Statement

D.A., G.M-R., S.K.S., P.W., L.R., J.S., T.M., T.T., B.G., C.M., A.M., S.G., J.F., J.H., M.J.M. and J.R.M are employed by Moderna Inc. and hold equities from the company.

